# Unmasking Anxiety in Autism: Explicit and Implicit Threatening Face Stimuli Dissociate Amygdala-centered Functional Connectivity

**DOI:** 10.1101/2020.03.24.005272

**Authors:** Yu-Chun Chen, Chenyi Chen, Róger Marcelo Martínez, Yang-Tang Fan, Chia-Chien Liu, Yawei Cheng

**Author notes:** Corresponding Authors: Yawei Cheng, M.D., Ph.D., Institute of Neuroscience and Brain Research Center, National Yang-Ming University, No. 155, Sec. 2, St. Linong, Dist. Beitou, Taipei 112, Taiwan, R.O.C. Equally contributed to the manuscript (Yu-Chun Chen, Chenyi Chen).

## Abstract

**Background:** Anxiety is the most prevalent comorbidity in individuals diagnosed with autism spectrum disorder (ASD). Amygdala reactivity to explicit and implicit threat processing is predictive of anxiety-related symptomatology. The neural mechanisms underlying the link between anxiety and ASD remains elusive.

**Methods:** In this fMRI study, we recruited young adults with ASD (N = 31) and matched them with controls, then proceeded to assess their autistic and anxiety traits by the use of the Autism-Spectrum Quotient (AQ) and the State-Trait Anxiety Inventory (STAI-S), respectively; and scanned their hemodynamic responses, including amygdala, in response to explicit and implicit (backwardly masked) perception of threatening faces.

**Results:** As compared to controls, the amygdala reactivity in ASD subjects was significantly lower to explicit threat, but comparable for implicit threat. The correlations of the amygdala reactivity with the AQ and STAI-S were dissociated depending on threat processing (explicit or implicit). Furthermore, the amygdala in ASD relative to controls had a more negative functional connectivity with the superior parietal cortex, fusiform gyrus, and hippocampus for explicit threat, whereas a more positive connectivity with the medial prefrontal cortex, temporal pole, and hippocampus for implicit threat.

**Conclusion:** In ASD, the transmission of socially relevant information along dorsal and ventral neural pathways centered on the amygdala is dissociated depending on explicit and implicit threat processing. This dissociation, ascribed to their failure to compromise pre-existing hyperarousal, might contribute to anxiety in ASD.

## BACKGROUND

Anxiety is the most prevalent co-occurring illness in individuals diagnosed with autism spectrum disorder (ASD)(Rodgers and Ofield, 2018). ASD is characterized by impaired social interaction, alongside stereotyped behavior and restrictive interests (APA, 2013). Previous conceptualizations regarding the underlying neuropathological changes central to the impairments in social interaction incurred by ASD have emphasized amygdala dysfunction (Amaral, et al., 2003; Baron-Cohen, et al., 2000; Schultz, 2005). Concurrently, the amygdala is equally central to the threat processing system involved in the experience of anxiety (Sah, 2017). However, the mechanisms underlying the link of anxiety and amygdala dysfunction with ASD remain elusive.

Whilst anxiety is a real and serious problem for many people on the autistic spectrum, relatively few neuroimaging studies have addressed this issue. One preliminary study with a small sample size (N = 12) reported that adults with ASD exhibited comparable amygdala engagement in response to implicit (backward masking) presentations of anxious (fearful) faces (Hall, et al., 2010). Another study used fear conditioning tasks (N = 20), showing anxiety was related to an impairment in the differentiation between threat versus safe cues in the amygdala (Top, et al., 2016). These findings suggest that ASD subjects might experience anxiety via a different route.

The amygdala, as a key brain structure implicated in anxiety, is responsible for saliency detection in the environment, including threats (Davis and Whalen, 2001). As such, the magnitude of amygdala reactivity in response to threatening stimuli (anger and fear) has been associated with anxiety (Etkin, et al., 2004; Etkin and Wager, 2007; Most, et al., 2006). Given that the threat-processing system is more reactive to ambiguous cues, the amygdala appears more reactive to fearful than angry faces (Whalen, et al., 2001), as fearful faces signify the presence of danger without providing information about its source. Consequently, amygdala reactivity to fearful faces is a reliable way of probing the neural correlates underlying anxiety (Bishop, et al., 2004; Chen, et al., 2017). Neuroimaging research has shown that the amygdala is engaged in implicit processing in addition to explicit processing of threats (Morris, et al., 1998; Whalen, et al., 1998). Implicit processing is presumed to be subserved by a direct neural pathway to the amygdala, thus permitting threat stimuli to be processed rapidly, automatically, and outside of conscious awareness (LeDoux, 1996). The paradigm using fMRI in conjunction with backwardly masked stimulus presentation provides a unique opportunity to examine behavioral and neural responses indicative of implicit threat processing (Dimberg, et al., 2000; Morris, et al., 2001; Whalen, et al., 1998). While non-masked stimuli for the explicit processing consisted of displaying fearful faces for 200-ms, backwardly masked stimuli for the implicit processing consisted of 17-ms of fearful faces followed by 183-ms of neutral faces.

Atypical attention to social stimuli in ASD has been well demonstrated (Fan, et al., 2014; Klin, et al., 2002). Moreover, heightened hemodynamic response in the amygdala has been associated with longer gaze fixation in autism (Dalton, et al., 2005). When explicitly perceiving socioemotional/threatening stimuli, individuals with ASD might tend to use attentional avoidance patterns to restrict affective hyperarousal. Conversely, and since implicit perception has been used to refer to a perceptual state in which the subjects do not report the presence of a stimulus even though the stimulus has in fact been processed (Tamietto and de Gelder, 2010), individuals with ASD might fail to inhibit pre-existing hyperarousal ascribed to anxiety when implicitly perceiving socioemotional/threatening stimuli.

Here, the fMRI study used the backwardly masked paradigm to elucidate how perceiving explicit and implicit threat affects the engagement of the amygdala and functional connectivity across two participant groups: ASD and controls. In regards to anxiety as indicated by amygdala reactivity to threat (Bishop, et al., 2004; Etkin, et al., 2004), we hypothesized that implicit anxiety in individuals with ASD might outweigh the explicitly induced fear. It is reasonable to suppose that anxiety in ASD might arise from the dissociation between amygdala reactivity to explicit and implicit threat processing. ASD could assert aberrant connectivity centered on the amygdala, compromising pre-existing hyperarousal triggered by implicit perception of threatening stimuli.

## MATERIALS AND METHODS

### Subjects

Thirty-one subjects with ASD and thirty matched controls participated in this study. Because of poor fMRI quality due to excessive head movement (a threshold of 1.75 mm, approximately half the size of a functional voxel, as the maximum head motion allowed in any plane), 29 subjects with ASD and 28 controls were finally included in the data analysis. The participants with ASD (aged 20.2 ± 6.0 years old, five female) were recruited from a community autism program. We reconfirmed the diagnosis of ASD by using the Diagnostic and Statistical Manual of Mental Disorders 5^th^ Edition’s (DSM-5) diagnostic criteria (APA, 2013). The participants in the age- and sex-matched control group (22.3 ± 3.5 years old, eight females) were recruited from the local community, and screened for major psychiatric illnesses by conducting structured interviews. The subjects did not participate in any intervention or drug programs during the experimental period. Participants with a comorbid psychiatric or medical condition, history of head injury, or genetic disorder associated with autism were excluded. All of them had normal or corrected-to-normal visual acuity. All participants, or their respective legal guardians, provided written informed consent All procedures were approved by the Ethics Committee and conducted in accordance with the Declaration of Helsinki.

### Procedures

Before fMRI scanning, each participant underwent assessments with the State-Trait Anxiety Inventory (STAI) to determine their self-reported anxiety levels (Spielberger, et al., 1970), as well as the Autism-Spectrum Quotient (AQ) (Baron-Cohen, et al., 2001b).

The paradigm for fMRI scanning was derived from the work by Etkin et al. (2004). The visual stimuli consisted of black and white pictures of male and female faces with fearful and neutral facial expressions, which were chosen from the Pictures of Facial Affect (Ekman and Friesen, 1976). The faces were oriented to maximize inter-stimulus alignment of eyes and mouths, and then artificially colorized (red, yellow, or blue) and equalized for luminosity. During fMRI scanning, subjects performed the color identification task, in which they were asked to judge the color of each face (pseudo-colored in either red, yellow, or blue) and to indicate the answer by a keypad button press. Each stimulus presentation involved a 200-ms fixation cross to cue subjects to focus on the center of the screen, followed by a 400-ms blank screen and a 200-ms face presentation. Participants then had 1200-ms to respond with a key press, indicating the color of the face. Non-masked stimuli consisted of a 200-ms fearful- or neutral-expression face. Backwardly masked stimuli consisted of 17-ms of a fearful or neutral face, followed by a 183-ms neutral face mask belonging to a different individual, but of the same color and gender as the previous one. Each epoch (12-s) consisted of six trials of the same stimulus type [Explicit Fearful (EF), Explicit Neutral (EN), Implicit Fearful (IF), or Implicit Neutral (IN)], but were randomized with respect to color and gender. The presentation order of the total 12 epochs (two for each stimulus type) and 12 fixation blocks (with a 12-s fixation cross) were pseudo-randomized. To avoid stimulus order effects, we used two different counterbalanced run orders. The stimuli were presented using Matlab software (MathWorks, Inc., Sherborn, MA, USA) and were triggered by the first radio frequency pulse for the functional run. The stimuli were displayed on VisuaStim XGA LCD screen goggles (Resonance Technology, Northridge, CA). The screen resolution was 800 x 600, with a refresh rate of 60 Hz. Behavioral responses were recorded by a fORP interface unit and saved in the Matlab program.

Immediately after fMRI scanning, participants underwent the detection task, during which they were shown all of the stimuli again and alerted of the presence of fearful faces. The subjects were administered a forced-choice test under the same presentation conditions as those during scanning and asked to indicate whether they observed a fearful face or not. The detection task was designed to assess possible awareness of the masked fearful faces. The chance level for correct answers was 50%. The performance was determined by the calculation of a detection sensitivity index (*d*) based on the percentage of trials in which a masked stimulus was detected when presented [‘hits’ (H)] and adjusted for the percentage of trials a masked stimulus was ‘detected’ when not presented [‘false alarms’ (FA)]; [*d’* = z-score (percentage H) − z-score (percentage FA), with chance performance = 0±1.74] (Kim, et al., 2010; Whalen, et al., 2004).

### Functional MRI data acquisition, image processing and analysis

Functional and structural MRI data were acquired on a 3T MRI scanner (Siemens Magnetom Tim Trio, Erlanger, German) equipped with a high-resolution 32-channel head array coil. A gradient-echo, T2*-weighted echoplanar imaging (EPI) with a blood oxygen level-dependent (BOLD) contrast pulse sequence was used for functional data. To optimize the BOLD response in the amygdala (Morawetz, et al., 2008), twenty-nine interleaved slices were acquired along the AC-PC plane, with a 96×128 matrix, 19.2×25.6 cm^2^ field of view (FOV) and 2×2×2 mm voxel size, resulting in a total of 144 volumes for the functional run (TR= 2-s, TE= 36-ms, flip angle=70°, slice thickness 2 mm, no gap). Parallel imaging GRAPPA with factor 2 was used to increase the speed of acquisition. Structural data were acquired using a magnetization-prepared rapid gradient echo sequence (TR= 2.53-s, TE= 3.03-ms, FOV= 256×224 mm^2^, flip angle= 7°, matrix= 224×256, voxel size= 1.0×1.0×1.0 mm^3^, 192 sagittal slices/slab, slice thickness= 1 mm, no gap).

Image processing and analysis were performed using SPM8 (Wellcome Department of Imaging Neuroscience, London, UK) in MATLAB 7.0 (MathWorks Inc., Sherborn, MA, USA). Structural scans were coregistered to the SPM8 T1 template, and a skull-stripped image was created from the segmented gray matter, white matter, and CSF images. These segmented images were combined to create a subject-specific brain template. EPI images were realigned and filtered (128-s cutoff), then co-registered to these brain templates, normalized to MNI space, and smoothed (4 mm FWHM). The voxel size used in the functional analysis was 2×2×2 mm^3^. All subjects who completed scanning had less than 1 voxel of in-plane motion.

Preprocessing for the T1-weighted images involved using the following DARTEL algorithm: new segment – generate roughly aligned gray matter (GM) and white matter (WM) images of the subjects; create template – determine nonlinear deformations for warping all the GM and WM images so that they match each other; and normalize to Montreal Neurological Institute (MNI) space – images were normalized to the MNI template and were smoothed with an 8-mm full-width at half-maximum Gaussian filter. Then, the GM, WM, and cerebrospinal fluid (CSF) structures of each patient were obtained after processing. A two-level approach for block-design fMRI data was adopted using general linear model implemented in SPM8. Fixed effects analyses were performed at the single subject level to generate individual contrast maps, and random effects analyses were performed at the group level. At single subject level, contrast images were calculated comparing each explicitly and implicitly presented fearful face block with the neutral baseline. Shorthand (e.g., EF−EN) was used to indicate the contrasts of regressors (e.g., Explicit Fearful > Explicit Neutral). Error bars signify SEM. To isolate the effects of fear content of the stimuli from other aspects of the stimuli and the task, we subtracted neutral (EN) or masked neutral (IN) activity from fearful (EF) or masked fearful activity (IF), respectively. The explicit perception of fearful faces was denoted as non-masked fear (EF−EN), and the implicit perception of fearful faces as masked fear (IF−IN). These contrast images were then entered to the second level group analysis. The resulting first-level contrast images were then entered into a full factorial analysis: 2 (group: ASD vs. CTL) x 2 (attention: explicit vs. implicit). Whole brain activations were corrected for multiple comparisons family-wise error (FWE) rate at *P*< .05.

Using MarsBar (see http://marsbar.sourceforge.net/), regions of interest (ROIs) analyses were conducted for the right and left amygdala according to prior meta-analyses (Costafreda, et al., 2008). Signals across all voxels with a radius of 5 mm were averaged and evaluated for the masked and non-masked comparisons in the right and left amygdala, respectively. The individual mean parameter estimates (beta values) were then subject to a mixed ANOVA, as to test for main effects of group (ASD vs. CTL) and attention (explicit vs. implicit), as well as group-by-attention interactions.

### Functional connectivity analysis

The psychophysiological Interaction (PPI) assesses the hypothesis that the activity in one brain region can be explained by an interaction between cognitive processes and hemodynamic activity in another brain region. The interaction between the first and second regressors represented the third regressor. The individual time series for the right amygdala was obtained by extracting the first principal component from all raw voxel time series in a sphere (5-mm radius) centered on the coordinates of the subject-specific amygdala activations. These time series were mean-corrected and high-pass filtered to remove low-frequency signal drifts. The physiological factor was then multiplied by the psychological factor to constitute the interaction term. PPI analyses were then carried out for each subject, and involved the creation of a design matrix with the interaction term, the psychological factor, and the physiological factor as regressors. PPI analyses were separately conducted for each group (ASD vs. CTL) in order to identify brain regions showing significant changes in functional coupling with the amygdala during explicitly and implicitly perceived fear. Subject-specific contrast images were then entered into random effects analyses to compare the group effect. Monte Carlo simulation implemented using AlphaSim (Ward, 2000) determined that a 5-voxel extent at height threshold of *P*< .005 uncorrected yielded a FWE corrected threshold of *P*< .05, accounting for spatial correlations in neighboring voxels. Subsequently, a multiple regression model was run separately for each seed to estimate the regression coefficient between all voxels and the interaction time series (along with task, movement, and linear drift as nuisance regressors). The strength of association between all voxels and the interaction time series was measured with R2 values. These coefficients of determination were square-rooted then multiplied by the sign of their respective estimated beta weights to obtain directionality of association. The correlation coefficients of the interaction time series were then converted to z-scores using Fisher’s transformation. The resulting statistical maps were then included in a second-level group analysis (ASD vs. CTL) by running a voxel-based two-sample *t*-test on the z-scores of the interaction effect for each seed separately.

## RESULTS

### Demographics and Dispositional Measures

The demographics and clinical variables of the participants are listed in Table 1. The ASD group, compared with the control group, scored higher on the AQ (*t*_38_= 4.07, *P* < .001, Cohen’s *d*= 1.1) in all of the subscales, as well as the STAI-T (*t*_38_= 2.37, *P* = .021, Cohen’s *d*= 0.6) and STAI-S (*t*_38_= 2.19, *P*= .032, Cohen’s *d*= 0.5).

**Table 1:**
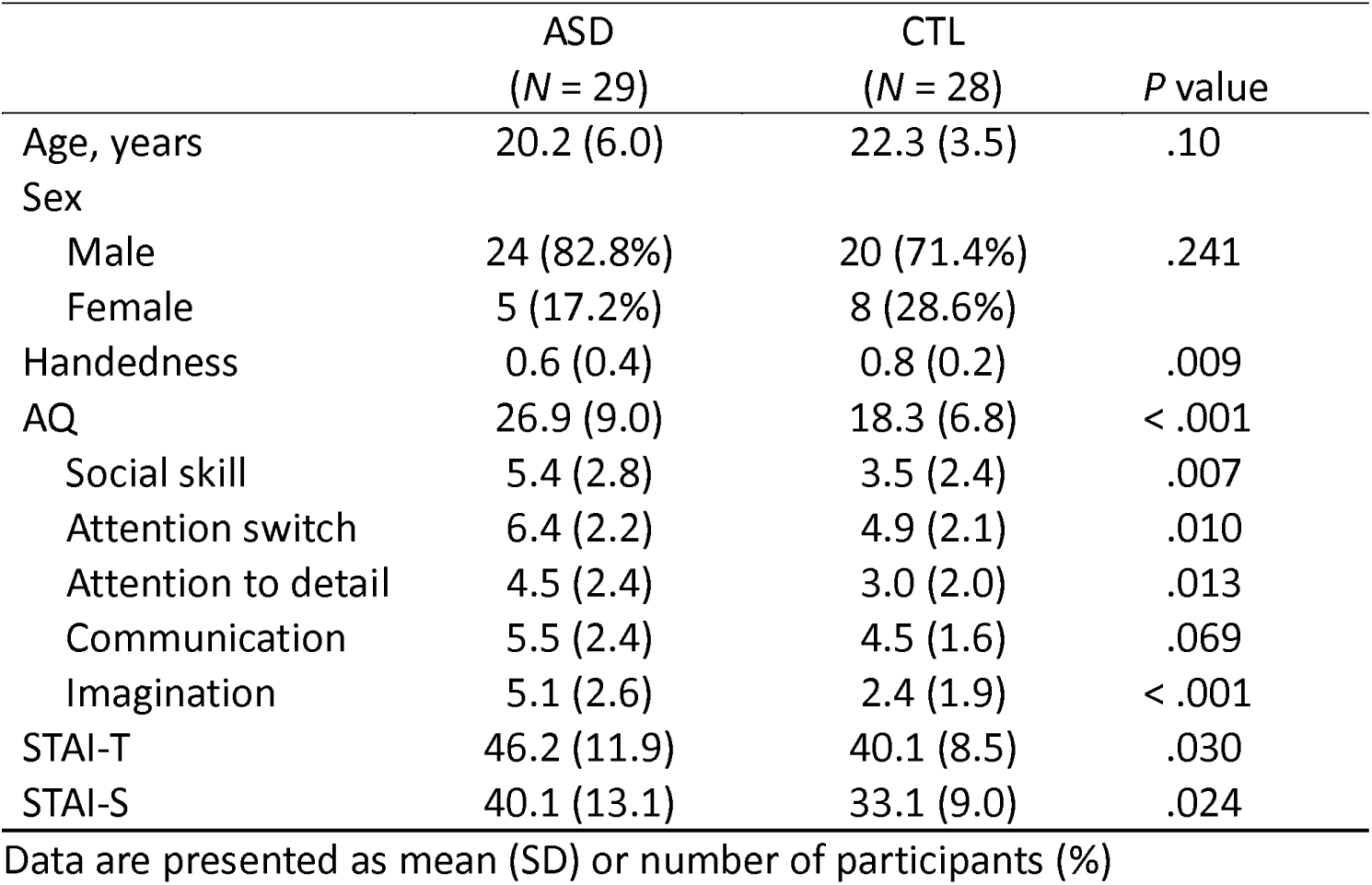
Demographic and clinical variables of the participants in the study.

|  | ASD<br>(N = 29) | CTL<br>(N = 28) | P value |
| --- | --- | --- | --- |
| Age, years | 20.2 (6.0) | 22.3 (3.5) | .10 |
| Sex |  |  |  |
| Male | 24 (82.8%) | 20 (71.4%) | .241 |
| Female | 5 (17.2%) | 8 (28.6%) |  |
| Handedness | 0.6 (0.4) | 0.8 (0.2) | .009 |
| AQ | 26.9 (9.0) | 18.3 (6.8) | < .001 |
| Social skill | 5.4 (2.8) | 3.5 (2.4) | .007 |
| Attention switch | 6.4 (2.2) | 4.9 (2.1) | .010 |
| Attention to detail | 4.5 (2.4) | 3.0 (2.0) | .013 |
| Communication | 5.5 (2.4) | 4.5 (1.6) | .069 |
| Imagination | 5.1 (2.6) | 2.4 (1.9) | < .001 |
| STAI-T | 46.2 (11.9) | 40.1 (8.5) | .030 |
| STAI-S | 40.1 (13.1) | 33.1 (9.0) | .024 |

Data are presented as mean (SD) or number of participants (%)

### Behavioral Performance

For the color identification task within the fMRI scanning, the two (group: ASD vs. CTL) x two (attention: explicit vs. implicit) x two (emotion: Fear vs. Neutral) ANOVA on the accuracy (± SEM) did not yield any effect (all *P* > .1). ASD individuals and controls appeared to have comparable performance (96.5 ± 4% vs. 96.1±5%). As regarding to the reaction time (RT), the results showed a main effect of emotion (*F*_1, 55_ = 11.0, *P* = .002, η^2^ = 0.17) and group (*F*_1, 55_ = 6.72, *P* = .012, η^2^ = 0.11) as well as an interaction between attention and emotion (*F*_1, 55_ = 4.11, *P* = .048, η^2^ = 0.07). While ASD and CTL did not differ in the accuracy, ASD took longer RTs in the color identification task (556±34 vs. 441±35 ms). While overall fear relative to neutral exerted longer RTs (514±26 vs. 493±23 ms), this negativity bias was unbalancedly biased towards the implicit condition (explicit: 506±26 vs. 493±24; implicit: 523±26 vs. 492±23).

For the detection task outside the fMRI scanner, according to a one-tailed binominal model, scores of 25 hits and above were considered significantly over chance level (50%). ASD and controls performed above chance level in the explicit condition (*d’*, mean ± SD: 2.084±0.941 and 2.162±0.780), and below chance level in the implicit condition (0.095±0.186 and 0.077±0.082). Between the groups, the overall accuracy (mean percentage of ‘hits’ and ‘correct rejection’) were not significantly different (explicit: *t*_55_ = −0.59, *P* = .557; implicit: *t*_55_ = 0.77, *P* = .442).

### Whole-brain fMRI results

The voxel-wise analysis identified the hemodynamic responses between groups in response to explicit and implicit fearful faces (Table 2). In response to explicit fear (EF−EN), as compared to the controls, the ASD subjects exhibited lower BOLD responses in the amygdala bilaterally, as well as in the left parahippocampus, orbitofrontal cortex (OFC), posterior cingulate cortex, and right precuneus [(EF−EN)|ASD < (EF−EN)|CTL]. For implicit fear (IF−IN), as compared to the controls, the BOLD response in the ASD subjects was higher in the right amygdala but lower in the left insula [(IF−IN)|ASD < (IF−IN)|CTL].

**Table 2:**
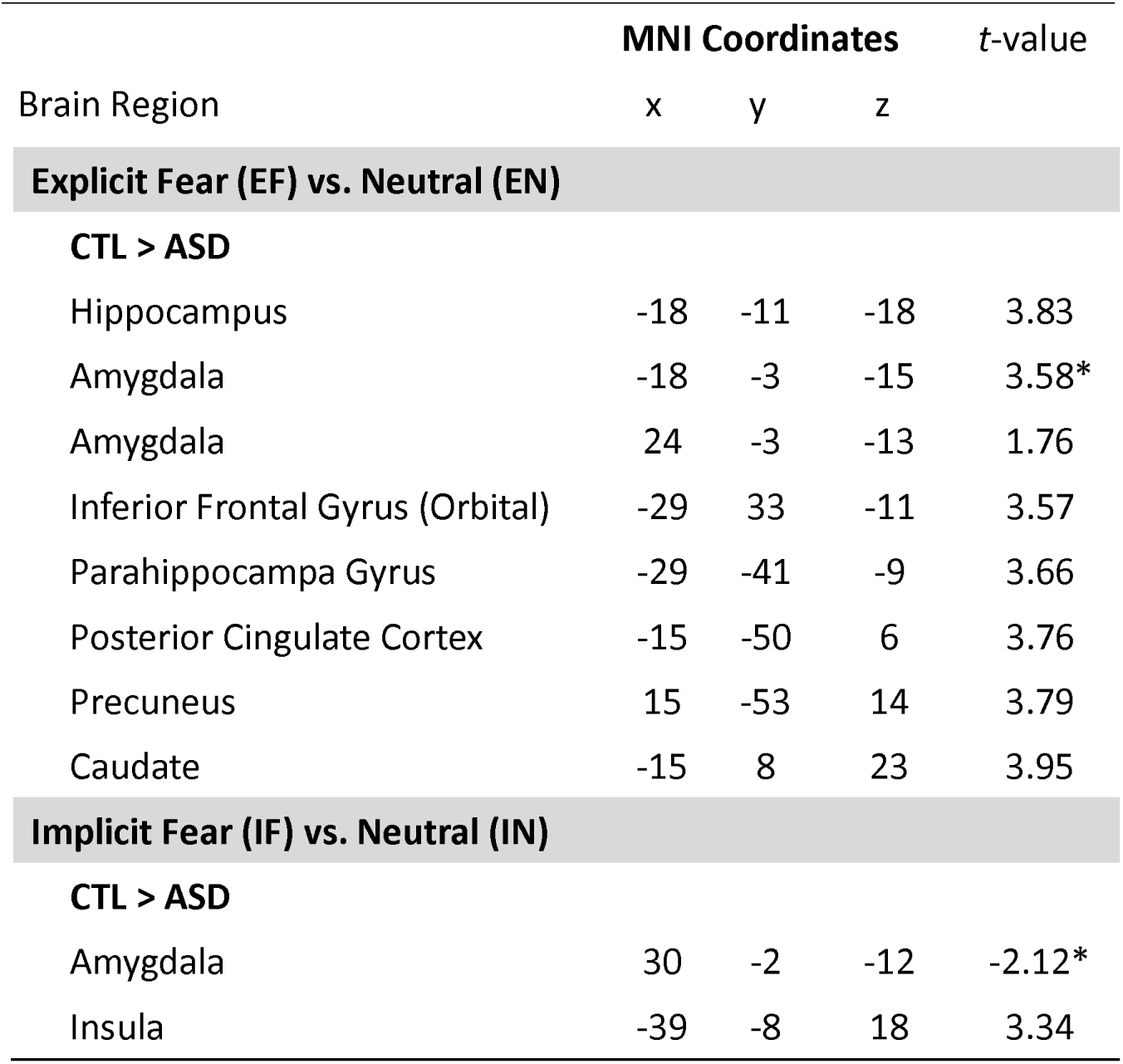
Group-wide fMRI results to explicit and implicit threat. All clusters are significant at FWE-corrected *P* < .05 [thresholded at *P* < .001, cut-off, *t* = 3.166 (uncorrected) with a spatial extent threshold *k* > 5], except those marked with an asterisk, which are taken from predefined ROIs and significant at uncorrected *P* < .05.

**Explicit Fear (EF) vs. Neutral (EN)****CTL > ASD**
|  |  |  |  |  |
| --- | --- | --- | --- | --- |
| Hippocampus | -18 | -11 | -18 | 3.83 |
| Amygdala | -18 | -3 | -15 | 3.58* |
| Amygdala | 24 | -3 | -13 | 1.76 |
| Inferior Frontal Gyrus (Orbital) | -29 | 33 | -11 | 3.57 |
| Parahippocampal Gyrus | -29 | -41 | -9 | 3.66 |
| Posterior Cingulate Cortex | -15 | -50 | 6 | 3.76 |
| Precuneus | 15 | -53 | 14 | 3.79 |
| Caudate | -15 | 8 | 23 | 3.95 |
| **Implicit Fear (IF) vs. Neutral (IN)****CTL > ASD** |  |  |  |  |
| Amygdala | 30 | -2 | -12 | -2.12* |
| Insula | -39 | -8 | 18 | 3.34 |

### Amygdala reactivity

To avoid circular inferences, prior ROI-based beta estimates were extracted from the amygdala bilaterally. In order to re-examine the validity of the backwardly masked paradigm, a priori comparison was conducted to inspect the simple main effect of fear versus neutral under the implicit condition in ASD and CTL, respectively. Results showed that fear relative to neutral showed significantly stronger right amygdala reactivity in CTL (*t*_27_ = 2.261, *P* = 0.032), but not in ASD (*t*_28_ = 0.942, *P* = 0.354). Subsequently, an ANOVA with one within-subject variable (explicit vs. implicit) and one between-subject variable (ASD vs. CTL) on the amygdala reactivity revealed interactions of group x attention (left: *F*_1, 55_ = 4.76, *P* = .033, η^2^ = 0.08; right: *F*_1, 55_ = 6.25, *P* = .015, η^2^ = 0.10). For the left amygdala, post hoc analyses indicated that ASD relative to controls showed significantly weaker amygdala reactivity to explicit fear (EF−EN: 0.06±0.155 vs. 0.553±0.075; *P* = .007), but they were comparable in response to implicit fear (IF−IN: −0.085±0.138 vs. −0.168±0.149; *P* = .687). For the right amygdala, post hoc analyses indicated that ASD relative to controls showed significantly weaker amygdala reactivity to explicit fear (0.027±0.116 vs. 0.30±0.060; *P* = .043), but they were comparable in implicit fear (0.134±0.157 vs. 0.064±0.076; *P* = .513) (Figure 1).

**Figure 1:**
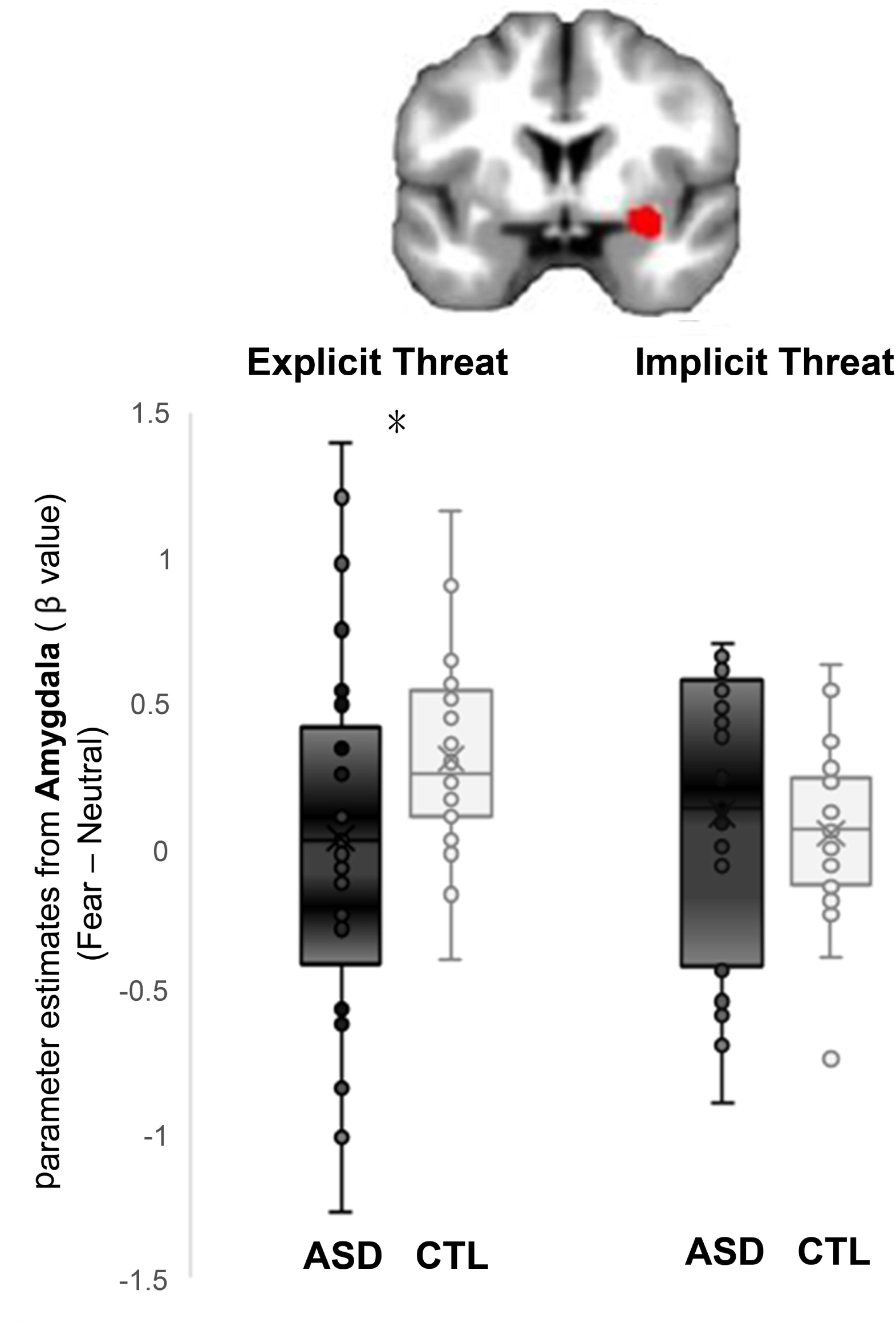
Dissociated amygdala reactivity between ASD and controls to explicit and implicit threat. The amygdala reactivity reveals an interaction of group (ASD vs. CTL) x attention (explicit vs. implicit) (*F*_1, 55_ = 6.25, *P* = .015). Post hoc analysis indicate that, as compared to controls, ASD individuals show significantly weaker amygdala reactivity in explicit fear (*P* = .043), but comparable in implicit fear (*P* = .513).

Furthermore, given that anxiety might involve a hyper-or-elevated baseline level of arousal to neutral stimuli (Canli and Lesch, 2007; Chen, et al., 2017; Top, et al., 2016), we further conducted detailed comparisons between explicit and implicit conditions to fearful and neutral faces (EF−IF; EN−IN), respectively. A two-way ANOVA with one within-subject variable (emotion: Neutral vs. Fearful) and one between-subject variable (group: ASD vs. CTL) was conducted for the right and left amygdala, respectively. In the right amygdala, there was an interaction of group x emotion (*F*_*1, 55*_ = 5.86, *P* = .019, η^2^ = 0.096). Post hoc analyses indicated that ASD relative to controls showed significantly stronger amygdala reactivity to neutral faces (EN−IN: 0.176±0.504 vs. −0.199±0.476; *P* = .006), but they were comparable in regards to fear (EF−IF: 0.155±0.618 vs. 0.163±0.271; *P* = .953). In the left amygdala, there was a marginal interaction of group x emotion (*F*_1, 55_ = 3.85, *P* = .055, η^2^ = 0.065). The effect of emotion in the amygdala had opposite directions depending on the factor of group. As compared to controls, ASD showed stronger amygdala activity when perceiving neutral faces whereas weaker amygdala activity during fear processing (Figure 2).

**Figure 2:**
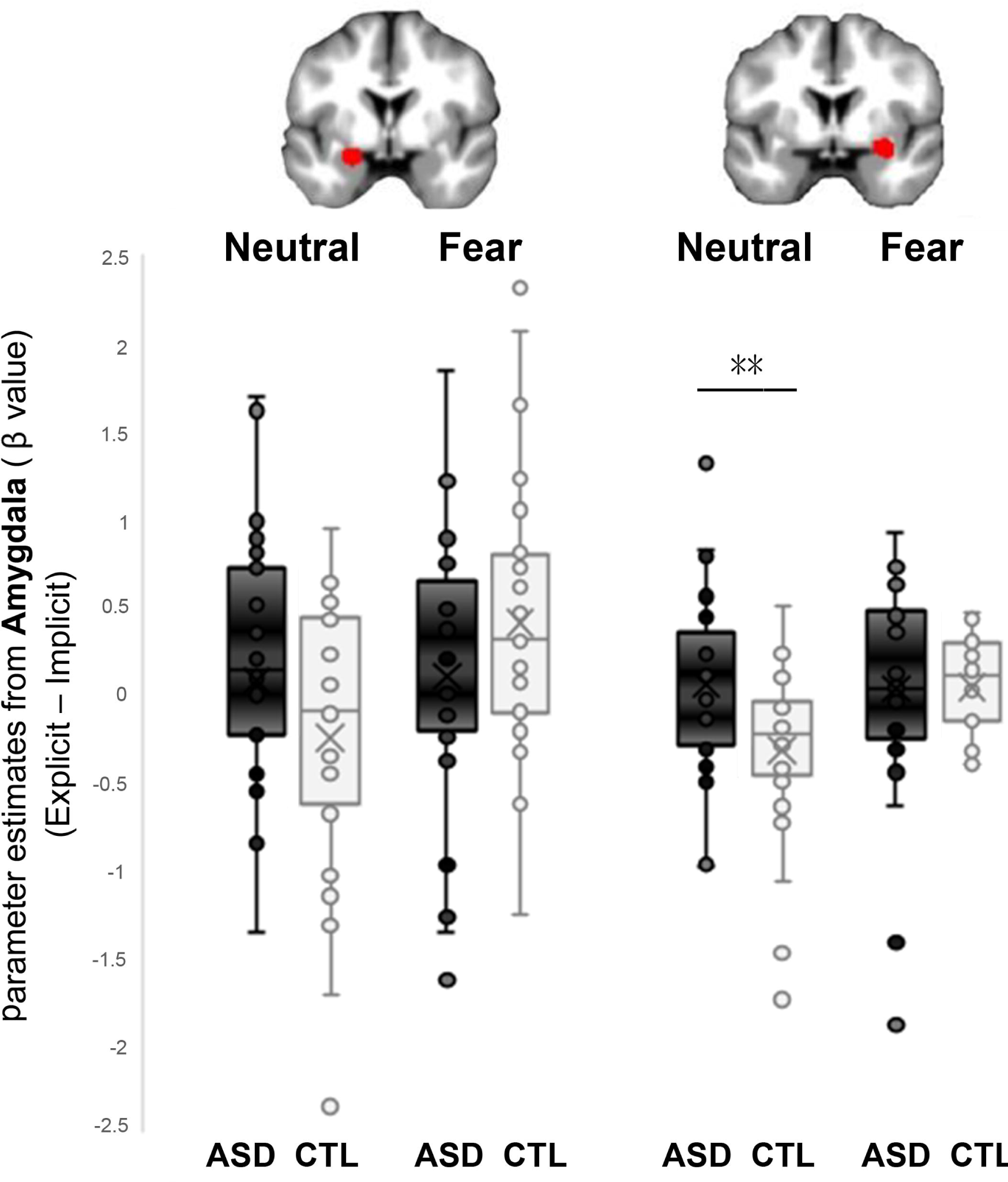
Over-reactivity to neutral faces in ASD. In the right amygdala, during the explicit vs. implicit condition, there is an interaction of group x emotion (*F*_1, 55_ = 5.86, *P* = .019). Post hoc analyses indicated that ASD relative to the controls showed significantly stronger amygdala reactivity to neutral faces (EN−IN: 0.176±0.504 vs. −0.199 ± 0.476; *P* = .006), but they were comparable in regards to fear (0.155±0.618 vs. 0.163±0.271; *P* = .953). In the left amygdala, there was a marginal interaction of group x emotion (*F*_1, 55_ = 3.85, *P* = .055). The effect of emotion in the amygdala has opposite directions depending on the factor of group. As compared to the controls, the ASD showed stronger amygdala activity to neutral faces but weaker amygdala activity to fearful faces.

To test if attention would dissociate the neural correlates of threat processing in ASD, we performed the Pearson product-moment correlation analysis between autistic trait and hemodynamic activity to explicit and implicit fear. When the ASD and control groups were recruited together (N= 57), the correlation between AQ and amygdala reactivity was negative in explicit fear (*r*= −0.37, *P*= .004), but positive in implicit fear (*r* = 0.33, *P* = .012) (Figure 3a). Fisher’s z tests confirmed a significant dissociation (z = −3.83, *P* < .001). Additionally, to test if attention would dissociate the neural correlates of threat processing related to anxiety, we performed the Pearson product-moment correlation analysis between STAI-S and amygdala reactivity to explicit and implicit fear. The correlation between STAI-S and amygdala reactivity was positive in implicit fear (*r* = 0.31, *P* = .026), but none in explicit fear (*r* = −0.08, *P* = .539) (Figure 3b). Fisher’s z tests confirmed a significant differential correlation (z = −2.04, *P* = 0.020).

**Figure 3:**
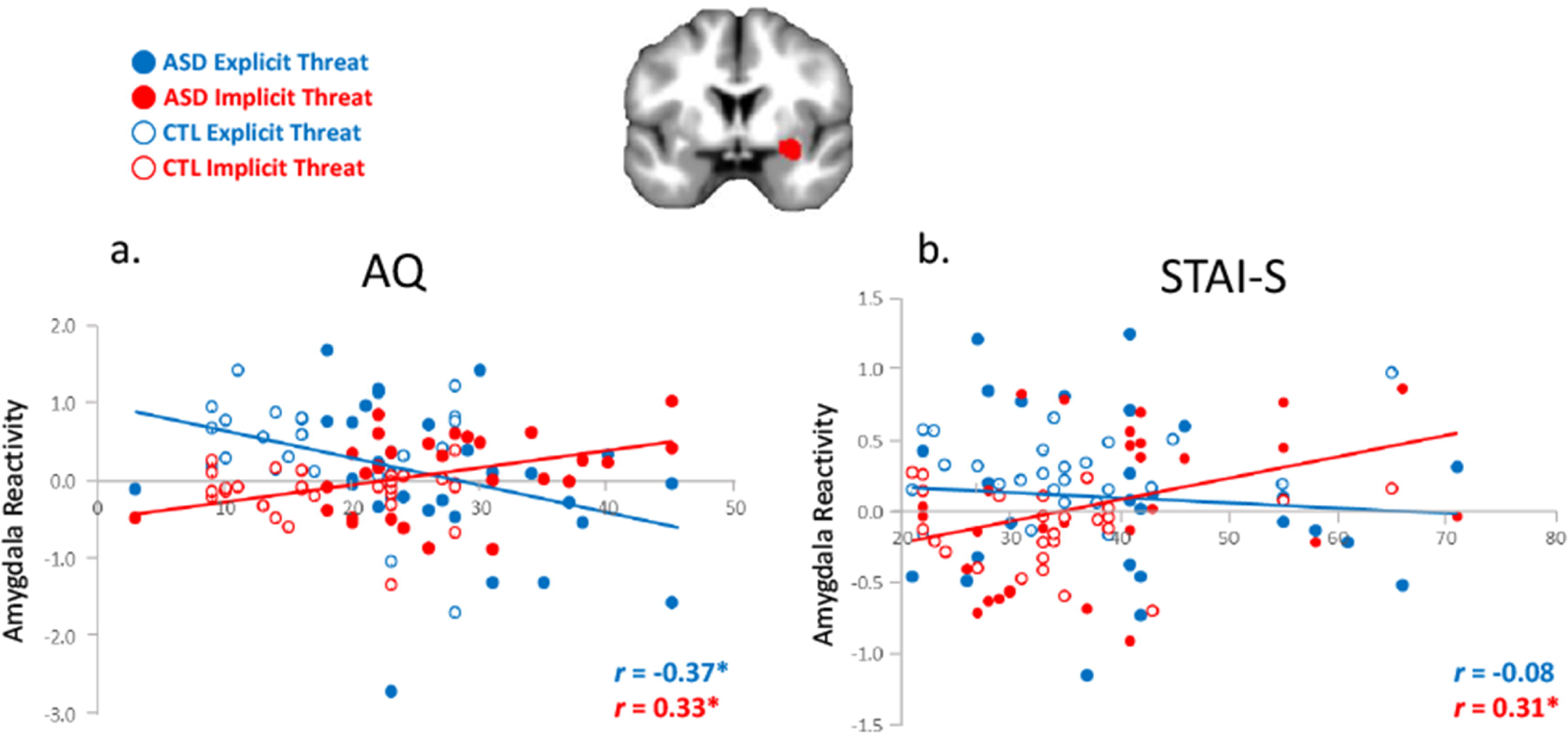
Dissociated correlations of autistic traits and anxiety with amygdala reactivity to explicit and implicit threat. **a**. The correlation between the autism quotient (AQ) and amygdala reactivity (30, 0, −26) is negative in explicit fear (*r* = −0.37, *P* = .004), but positive in implicit fear (*r* = 0.33, *P* = .012) when the ASD and control groups are recruited together (N = 57). Fisher’s *z* test confirms a significant dissociation (z = −3.83, *P* < .001). **b**. The correlation between STAI-S and amygdala reactivity was positive in implicit fear (*r* = 0.31, *P* = .026), but none in implicit fear (*r* = −0.08, *P* = .539). Fisher’s z tests confirmed a significant differential correlation (z = −2.04, *P* = .020).

### Functional Connectivity

To further examine the extent to which ASD-related modulation in threat processing contributed to the functional coupling between different brain regions, we subsequently assessed functional connectivity in the amygdala, whose functions are related to fear processing (Figure 4). The time series of the first eigenvariates of the BOLD response were temporally filtered, mean corrected, and deconvolved to generate the time series of the neuronal signal for the source region –the left and right amygdala (−18, −3, −15; 30, −2, −12)– as the physiological variable in PPI analysis. Being selected as the PPI source region, the physiological regressor was denoted by the activity in the amygdala. Fear was the psychological regressor. For explicit fear (EF−EN), as compared to the controls, the ASD showed a significantly more negative connectivity of the amygdala with the superior parietal cortex (−26, −64, 49), fusiform gyrus (38, −60, −14), and hippocampus (38, −22, −8). To further examine whether this effect was driven by a weaker positive coupling or stronger negative coupling in ASD, PPI analyses were conducted for ASD and controls separately for descriptive purposes (Table 3). There were aberrant negative couplings of the amygdala with the ventral and dorsal neural pathways when ASD perceived explicit threat. For implicit fear (IF−IN), as compared to the controls, the ASD showed a significantly stronger correlation of the amygdala with the medial prefrontal cortex (9, 59, −2), the temporal pole (45, 8, −18) and hippocampus (36, −16, −12).

**Table 3:** Group-wise fMRI results of the functional connectivity during implicit and explicit threat processing.

| Region | MNI coordinates |  |  | Cluster size | T |
| --- | --- | --- | --- | --- | --- |
|  | x | y | z |  |  |
| Explicit Threat |  |  |  |  |  |
| ASD |  |  |  |  |  |
| Fusiform Gyrus | 34 | -55 | -9 | 16 | -2.74<sup>†</sup> |
| Hippocampus | 38 | -22 | -8 | 82 | -3.28<sup>‡</sup> |
| Superior Parietal Cortex | -26 | -64 | 49 | 63 | -2.55<sup>†</sup> |
| CTL |  |  |  |  |  |
| Fusiform Gyrus | 34 | -55 | -9 | N.S. | N.S. |
| Hippocampus | 38 | -22 | -8 | N.S. | N.S. |
| Superior Parietal Cortex | -26 | -64 | 49 | N.S. | N.S. |
| Implicit Threat |  |  |  |  |  |
| ASD |  |  |  |  |  |
| Temporal Pole | 45 | 8 | -18 | 18 | 2.54<sup>‡</sup> |
| Hippocampus | 36 | -16 | -12 | 26 | 2.42<sup>†</sup> |
| Inferior Frontal Cortex | 51 | 33 | -9 | 12 | 2.41<sup>†</sup> |
| Medial Prefrontal Cortex | 9 | 59 | -2 | N.S. | N.S. |
| CTL |  |  |  |  |  |
| Temporal Pole | 45 | 8 | -18 | N.S. | N.S. |
| Hippocampus | 36 | -16 | -12 | N.S. | N.S. |
| Inferior Frontal Cortex | 51 | 33 | -9 | N.S. | N.S. |
| Medial Prefrontal Cortex | 9 | 59 | -2 | 14 | -3.03<sup>‡</sup> |

‡, $P = .006$ ; †, $P = .010$ .

**Figure 4:**
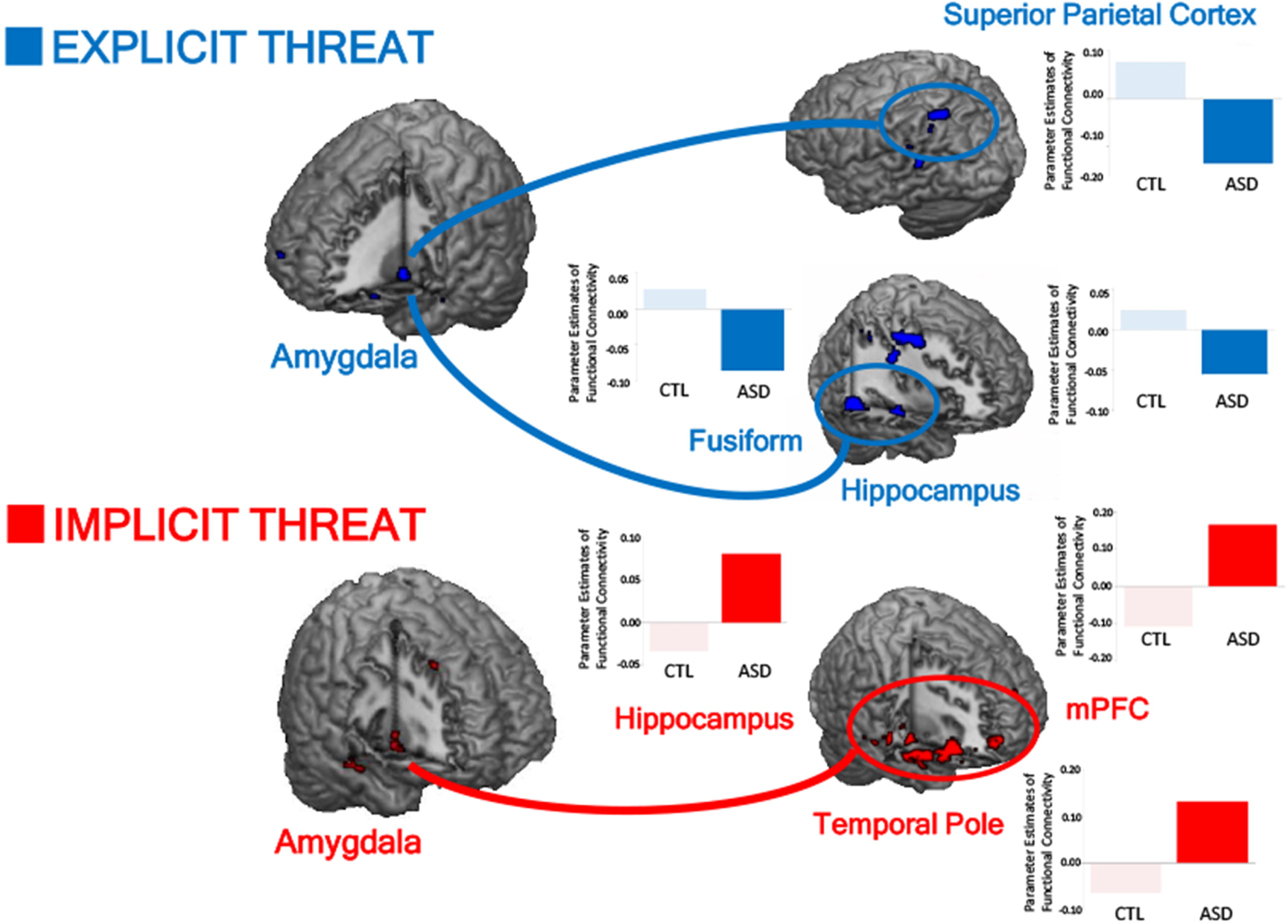
Dissociated amygdala functional connectivity by explicit and implicit threat in ASD. Compared to the controls, ASD subjects have a significantly more negative connectivity of the amygdala with the superior parietal cortex, fusiform gyrus, and hippocampus when processing explicit threat, whereas a significantly more positive connectivity of the amygdala with the medial prefrontal cortex (mPFC), temporal pole and hippocampus when processing implicit threat.

## DISCUSSION

It is important to understand the neural mechanisms underlying the link of anxiety and amygdala dysfunction in ASD. Here, we combined explicit and implicit (backward masking) perception of fearful faces to elicit amygdala reactivity in controls and participants with ASD, who varied in autistic trait and self-reported anxiety. The results showed that, as compared to the controls, amygdala reactivity was significantly lower towards explicit fear, but was comparable to implicit fear in ASD. The correlations of amygdala reactivity with the AQ and STAI-S were dissociated between explicit and implicit fear. Furthermore, in ASD relative to controls, the amygdala exhibited an aberrant and more negative functional connectivity with the superior parietal cortex, fusiform gyrus, and hippocampus in explicit fear, whereas a more positive connectivity with the medial prefrontal cortex, temporal pole, and hippocampus during implicit fear. On one hand, the existing over-reactivity to neutral faces, along with stronger negative couplings in both dorsal and ventral neural pathways during explicit threat, may explain why the failure to compensate the preexisting hyperarousal –even with the extra cost of energy consumption– in ASD would lead to a higher level of anxiety and more severe autistic traits (Top, et al., 2016). On the other hand, during implicit threat, the stronger positive connectivity along the ventral neural pathways indicates consciousness-dependent transmission of socially relevant information in ASD (Pessoa and Adolphs, 2010; Tamietto and de Gelder, 2010).

ASD shows weaker amygdala reactivity towards explicit fear, although its activity appears to be comparable to the controls during implicit fear. One preliminary study in adults with ASD uncovered heightened amygdala activation, but this research was limited to a backward masking paradigm with subthreshold presentations of fearful faces (Hall, et al., 2010). The present findings extend the existing scientific literature by demonstrating a significant dissociation dependent on explicit and implicit threat. Such a dissociation appears parallel to the empathy imbalance hypothesis of autism in regards to the surplus of emotional empathy in people with ASD (Smith, 2009). Compelling evidence concerning this excess in emotional empathy characteristic of autism comes from self-report questionnaires, facial electromyography, and event-related potentials. As shown by the Interpersonal Reactivity Index, which includes items designed to tap each facet of empathy, people with Asperger syndrome scored significantly higher than the controls on the personal distress scale (Rogers, et al., 2007). These findings indicate that people with Asperger’s syndrome might have a tendency to be anxious, and that this may contribute to a particular susceptibility towards empathic overarousal. As indicated by facial electromyography, which measures automatic mimicry of subtle emotional responses, youths with ASD showed significantly heightened electromyographic responsiveness when being presented with pictures of happy and fearful faces, and instructed to judge the gender of each face (Magnée, et al., 2007). This result contrasts with those of another study that found deficits in the automatic mimicry of emotional faces (McIntosh, et al., 2006). The crucial difference between the two studies lies on whether the subjects were set to an active task requiring careful attention to the faces or not. Deficits in eye contact, as indicated by aberrant visual attention, have been a hallmark of autism (Jones and Klin, 2013). In parallel, as shown by the event-related potentials for pain empathy, an early automatic component (N2) indexing empathic arousal, was heightened in individuals with ASD (Fan, et al., 2014). These findings are consistent with the hypothesis that the underlying capacity for emotional empathy in autism is not just intact, but excessive. People with autism may not actively attend to socioemotional information in order to minimize empathic hyperarousal.

The correlation between AQ and amygdala reactivity is dissociated, as it is dependent on whether threat is explicit or implicit. Among the variety of screening tools developed to quantify autistic traits, the most commonly used is probably the Autism-Spectrum Quotient (AQ)(Baron-Cohen, et al., 2001b). The AQ has been used to screen clinical samples (Woodbury-Smith, et al., 2005) and to predict performance on cognitive tasks (Stewart, et al., 2009), social cognition (Baron-Cohen, et al., 2001a), facial mimicry (Hermans, et al., 2009), gaze preference to social stimuli (Bayliss and Tipper, 2005), and speech perception (Stewart and Ota, 2008). In the same line, more severe autistic traits, as assessed by the AQ, were coupled with weaker mismatch negativity (MMN) (Fan and Cheng, 2014). Not until recently was the MMN, a component of the event-related potentials to an odd stimulus in a sequence of stimuli, utilized as an index for pre-attentive salience detection of emotional voice processing (Cheng, et al., 2012; Hung, et al., 2013; Hung and Cheng, 2014). The unexpected presence of emotional spoken syllables embedded in a passive oddball paradigm were able to activate the amygdala, which was associated with individual differences in social orientation (Schirmer, et al., 2008). The amygdala reactivity to explicit and implicit fearful faces exhibited opposite associations with MMN (Chen, et al., 2017). Thus, it is no surprise to see a dissociation regarding the coupling between AQ and amygdala reactivity dependent on explicit and implicit threat.

The amygdala-centered functional connectivity between explicit and implicit threat is dissociated in ASD. Although many studies have reported that individuals with ASD have atypical brain connectivity patterns, the results of more recent studies do not support unanimously the traditional view where individuals with ASD are described as having lower connectivity between distal brain regions and increased connectivity within proximal brain regions (Mohammad-Rezazadeh, et al., 2016). For instance, literature reviews observed a general trend supporting the hypothesis of long-range functional under-connectivity, however, the status of local connectivity remains unclear (O’Reilly, et al., 2017). Thus, further investigations of connectivity with respect to behavior are needed to probe the underlying brain networks implicated in core deficits of ASD. In the same vein, thalamocortical disconnectivity could account for sensorimotor symptoms in ASD (Woodward, et al., 2017). Restricted interest and repetitive behaviors could be determined via inter- and intra-hemispheric functional disconnectivity (Lee, et al., 2016). Here, this study reidentifies the dorsal and ventral networks as showing shared activity, and more importantly demonstrates the existence of attention dissociated amygdala-centered pathways, towards threat (Cole, et al., 2016; Friston, 2011). ASD showed significantly stronger negative signal couplings in both dorsal and ventral neural pathways during explicit threat processing, whereas stronger positive correlation along the ventral neural pathways during implicit threat processing.

## Limitations

Some limitation of this study must be acknowledged. First, regarding sample homogeneity, the generalizability of the results may be limited because people with low-functioning ASD were not included. Second, the relatively wide age range of subjects here might interact with developmental brain changes. Accordingly, based on the same theorems as previous work (Schipul and Just, 2016), we have conducted the imaging processing with DARTEL, as well as the correlation analysis with age and the inclusion of age as a covariate of no interest, as well as DARTEL corrections. This may not be the optimal design, and future studies are warranted with a larger sample size.

## Conclusion

Taken together, our study unmasks the nature of anxiety in ASD, as shown by the dissociation of amygdala reactivity and functional connectivity dependent on explicit and implicit threat processing. Implicit anxiety in individuals with ASD could outweigh the explicitly induced threat. When explicitly perceiving socioemotional stimuli, individuals with ASD might use attentional avoidance patterns to restrict affective hyperarousal. This gives a sense of urgency for the need to develop an attention-independent neural marker concerning anxiety in ASD.

## ACKNOWLEDGMENTS

We deeply thank the participants and their parents who were included in the study. The study was funded by the Ministry of Science and Technology (MOST 108-2410-H-010-005-MY3; 108-2636-H-038-001-; 108-2410-H-009 −020 -MY3; 109-2636-H-038-001-), National Yang-Ming University Hospital (RD2019-003; 2020-003).

## Author Contributions

Y.C.C., C.C., and Y.C. conceived and conceptualized the study. Y.C.C., Y.T.F., and C.C. collected and analyzed the data. Y.C.C., C.C., and Y.C. conducted the necessary literature reviews and drafted the first manuscript. R.M.M. and Y.T.F. provided critical feedback and helped shape the manuscript. All authors contributed towards the revision and writing the final draft.

