## Supplemental Figure 1 for "Unmasking Anxiety in Autism: Explicit and Implicit Threatening Face Stimuli Dissociate Amygdala-centered Functional Connectivity"

**SUPPLEMENTARY MATERIALS**

**Figure s1: fMRI results to explicit and implicit threat.**

**a.** In response to explicit fear (EF−EN), as compared to the controls, the ASD subjects exhibited lower BOLD responses in the amygdala bilaterally, as well as in the left parahippocampus, orbitofrontal cortex (OFC), posterior cingulate cortex, and right precuneus.

**b.** For implicit fear (IF−IN), as compared to the controls, the BOLD response in the ASD subjects was higher in the right amygdala but lower in the left insula.

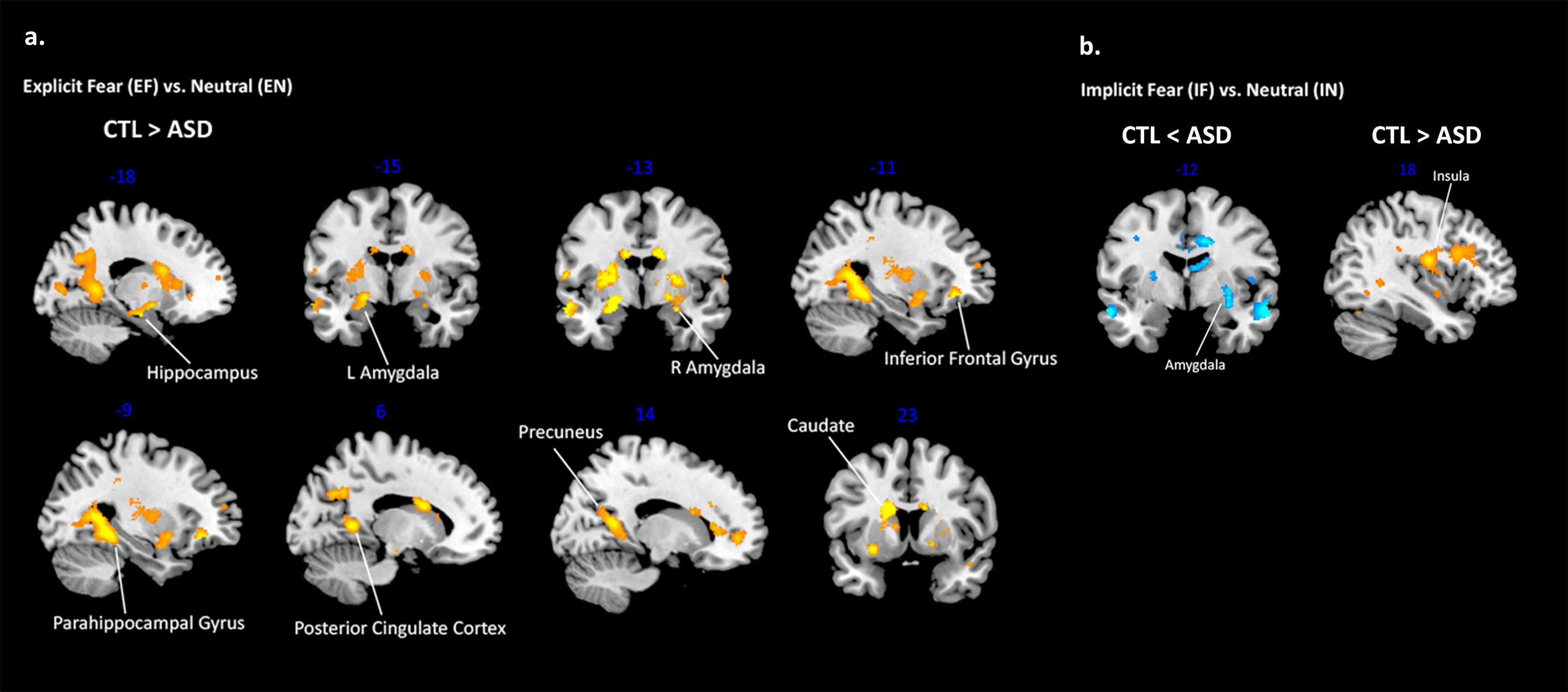
